# hictk: blazing fast toolkit to work with .hic and .cool files

**DOI:** 10.1101/2023.11.26.568707

**Authors:** Roberto Rossini, Jonas Paulsen

## Abstract

**Motivation:** Hi-C is gaining prominence as a method for mapping genome organization. With declining sequencing costs and a growing demand for higher-resolution data, efficient tools for processing Hi-C datasets at different resolutions are crucial. Over the past decade, the .hic and Cooler file formats have become the de-facto standard to store interaction matrices produced by Hi-C experiments in binary format. Interoperability issues make it unnecessarily difficult to convert between the two formats and to develop applications that can process each format natively.

**Results:** We developed hictk, a toolkit that can transparently operate on .hic and .cool files with excellent performance. The toolkit is written in C++ and consists of a C++ library with Python and R bindings as well as CLI tools to perform common operations directly from the shell, including converting between .hic and .mcool formats. We benchmark the performance of hictk and compare it with other popular tools and libraries. We conclude that hictk significantly outperforms existing tools while providing the flexibility of natively working with both file formats without code duplication.

**Availability:** The hictk library, Python bindings and CLI tools are released under the MIT license as a multi-platform application available at github.com/paulsengroup/hictk. Pre-built binaries for Linux and macOS are available on bioconda. Python bindings for hictk are available on GitHub at github.com/paulsengroup/hictkpy, while R bindings are available on GitHub at github.com/paulsengroup/hictkR.

**Supplementary information:** Supplementary data are available at *Bioinformatics* online.

## 1 Introduction

Hi-C and related tools involve probing and quantifying pairwise proximities throughout entire genomes’ three-dimensional (3D) structure, and are invaluable for studying genome- and chromatin organization. Hi-C data production has seen a dramatic rise recently, both for advancing genome assembly (Burton *et al*., 2013; Kaplan and Dekker, 2013; Selvaraj *et al*., 2013) and for precise mapping of the genome’s 3D architecture (Lieberman-Aiden *et al*., 2009), putting to test the performance of available computational tools to process and analyze Hi-C data.

Sequencing data produced by Hi-C experiments are usually processed with specialized pipelines, such as those provided by ENCODE, 4DNucleome, nf-core, and Open2C. The main file output by these pipelines stores a matrix-representation of all pairwise Hi-C interactions within and between the reference chromosomes.

These interaction matrices are typically very large (9.5 trillion values at 1 kbp resolution in the human genome) and thus cannot be represented as dense matrices in plain-text format.

Two popular file formats are currently used to efficiently store this kind of data: Cooler (Abdennur and Mirny, 2019) and .hic (Durand, Robinson, *et al*., 2016). Both formats represent interaction matrices as sparse, compressed and chunked matrices, yet in substantially different ways, requiring specialized libraries for reading or writing each format. As a consequence, application developers need to write two versions of the same code to read data from files: one for .hic files and another for Cooler files. Yet, responsibility of format conversion is usually placed on the users of the tools. Notably, conversion between the two formats is not straightforward or computationally efficient, requiring an intermediate text representation for reliable conversion. This process is inefficient and error prone, as users are tasked with verifying the accuracy of the text format and ensuring proper sorting.

Files stored in the .hic format undergo processing through several tools: JuiceboxGUI (Durand, Robinson, *et al*., 2016) offers an interactive interface where users can visualize .hic matrices at different resolutions, with various normalizations and annotation tracks. JuicerTools and HiCTools (Durand, Shamim, *et al*., 2016) facilitate the generation of .hic files, and straw (Durand, Robinson, *et al*., 2016) functions as a versatile .hic file reader. Notably, straw is accessible as a library for multiple programming languages such as C++, Matlab, R, and Python. In contrast, JuiceboxGUI and HiCTools are Java-based applications designed to efficiently handle .hic files.

Cooler files on the other hand, are processed with the homonymous suite of tools and can be visualized using HiGlass (Abdennur and Mirny, 2019; Kerpedjiev *et al*., 2018). The Cooler file format is based on HDF5 (www.hdfgroup.org/HDF5), with a CLI and library written in Python. Usage of HDF5 represents a difference in philosophy from the .hic format, allowing flexible and relatively user-friendly runtime-based introspection of data. However, users willing to open cooler files using programming languages other than Python still need to create dedicated code to e.g. satisfy queries by genomic coordinates. The Cooler file format specification defines three formats: single-resolution (.cool), multi-resolution (.mcool) and single-cell (.scool) Cooler files. Single-resolution Cooler files store interactions using a compressed sparse row (CSR) scheme; multi-resolution Coolers are collections of .cool files at different resolutions; single-cell Cooler files are meant to store single-cell Hi-C data. Contrary to files in Cooler format, files in .hic format can be visualized using the UCSC genome browser.

Finally, a tool named hic2cool is available to easily convert from .hic to .mcool. hic2cool is written in Python and borrows code from straw and cooler. However, hic2cool is relatively slow (see benchmarks) and is not capable of reading the latest version of .hic files (github.com/aidenlab/hic-format).

We have developed the hictk toolkit with the aim of streamlining reading, writing and converting .hic and Cooler files within the same library, and doing so with excellent performance. hictk consists of a library and CLI components. The library allows reading and writing both .hic and Cooler files. The CLI makes it easy to perform common operations directly from the shell, including converting files between Cooler and .hic formats (see Supplementary Fig. 1).

**Fig. 1:**
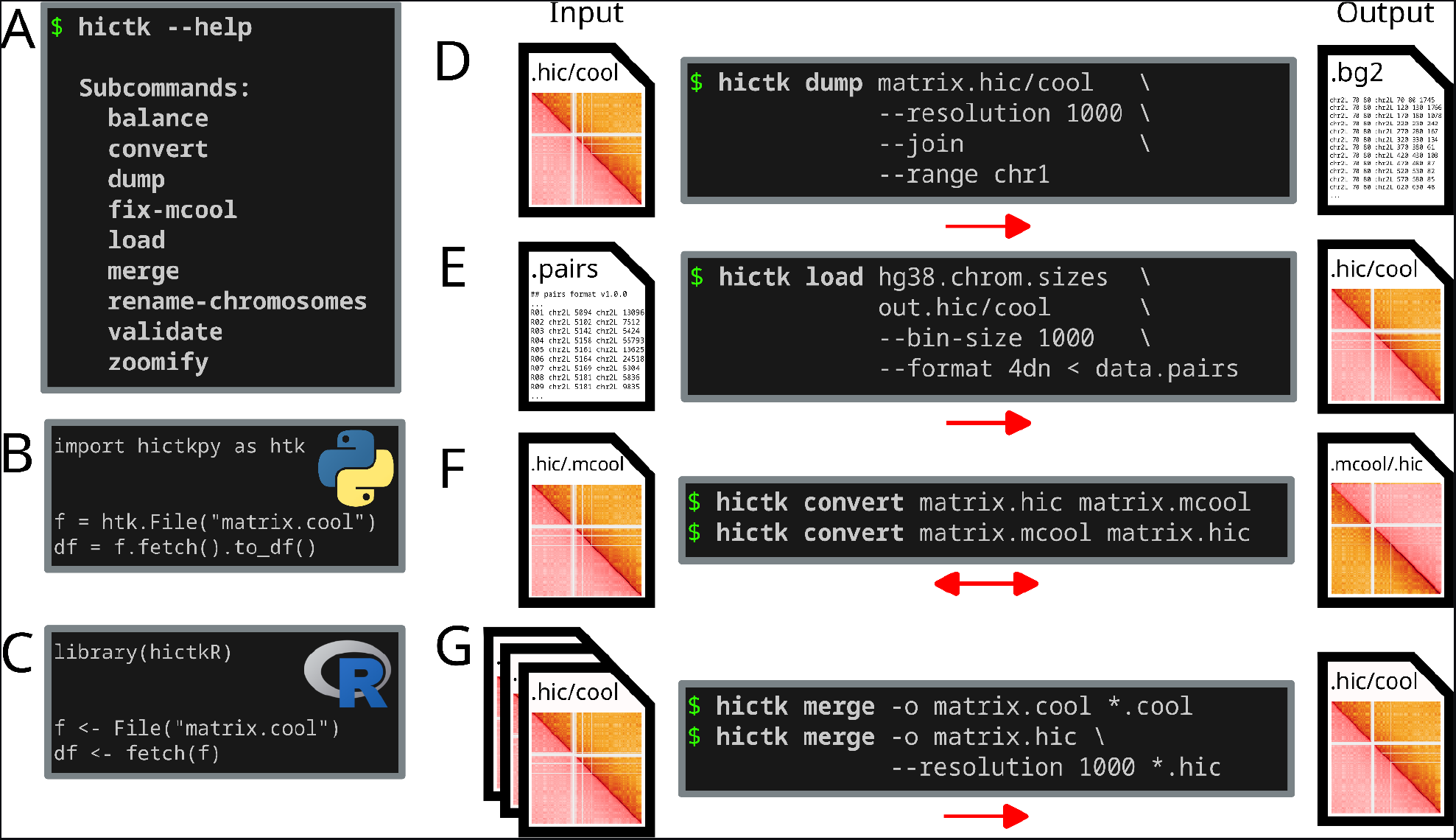
Overview of hictk functionality. **A** List of CLI subcommands provided by hictk. **B** Snipped of Python code showing how to use hictkpy to fetch genome-wide interactions from a .cool file. **C** Snippet of R code showing how to use hictkR to fetch genome-wide interactions from a .cool file. **D** Example usage of hictk dump. The example shows how to use the dump subcommand to extract chr1 interactions at 1000 bp resolution from a .hic or .cool file and print them to stdout as a list of bedGraph2 records. **E** Example usage of hictk load. The example shows how to ingest interactions in 4DN pairs format to generate a single-resolution .hic or .cool file. **F** Example usage of hictk convert showing how to convert a file in .hic format to .mcool and vice-versa. **G** Example usage of hictk merge showing how to merge multiple .hic or .cool files, producing a single file in the same format as the input files.

## 2 Material and methods

### 2.1 Overview of the hictk library

hictk is implemented in C++ and was designed with computational- and memory efficiency and composability in mind. To achieve this, hictk heavily relies on iterators to lazily traverse collections of pixels. From the user perspective, there is no difference between reading interactions from a .hic, .cool or .mcool file. To read interactions for a region of interest, users first create a file object by providing the path to a file and the resolution in bp to be used. The file object implements a fetch method that takes as input several optional parameters, including a query range (e.g. chr1:0-10,000,000). The fetch method returns a PixelSelector object, providing begin() and end() methods allowing pixel traversal for the queried range (see Supplementary Table 1 found in Supplementary Text 1). PixelSelector objects also support returning interaction frequencies directly as sparse or dense matrices. Iterators are implemented to strike a good balance between memory usage and performance, requiring few KBs of memory when fetching small regions on coarse matrices up to hundreds of MBs when fetching large queries from high resolution matrices while traversing millions of pixels per second (see benchmarks for more details).

The library supports reading files with integer and floating-point interaction frequencies, with and without matrix balancing using algorithms such as iterative correction and eigenvector decomposition (ICE) or Knight-Ruiz (KR) (Imakaev *et al*., 2012; Knight and Ruiz, 2012). For .hic files, expected and observed/expected interaction frequencies are also supported. This level of flexibility is achieved through runtime polymorphism enabled by the visitor pattern (Palsberg and Jay, 1998). Beside reading interactions, the library provides a set of operations that can be used as building blocks for more complex applications, such as functions to create .cool, .mcool and .scool files, merge multiple files, coarsening interactions from .hic and .cool files and more.

Unrecoverable errors are propagated through exceptions, letting the caller decide whether an application using hictk should abort in case of errors.

The API of hictkpy, hictk’s Python bindings, was heavily inspired by cooler’s Python API.

Beside performance, hictk’s C++ implementation allows writing bindings for popular scripting languages such as Python and R through e.g. nanobind github.com/wjakob/nanobind and Rcpp (Eddelbuettel, 2013). We currently provide bindings for the Python and R languages through the hictkpy and hictkR packages, respectively. In Section 9 in the Supplementary Text 1 we show how to update a Python application to use hictkpy instead of cooler to read contact matrices, making the application capable of reading .hic files with few changes.

### 2.3 Testing

hictk is thoroughly tested with a continuous integration pipeline (CI) which automatically executes hictk unit tests (80%+ coverage) on Linux, macOS and Windows using different versions of GCC, Clang, Apple-Clang and MSVC.

CLI tools are further tested with 20 integration tests, covering the most common use-cases of each subcommand.

As an example, hictk convert is tested by two different integration tests covering conversion from .mcool to .hic and vice-versa. After performing the conversion, the tests dump interactions for the original and converted files to two different text files. The text files are then compared and an error is raised in case differences are found.

Finally, fetch functionality is automatically tested every time a pull request is merged, or weekly, with thousands of random queries to ensure that hictk and cooler return identical results for the same query.

### 2.4 Documentation

Documentation for hictk’s C++ API and CLI interface is available on ReadTheDocs at the following address: hictk.readthedocs.io. The documentation also contains instructions on how to install or build hictk. Documentation for hictkpy and hictkR are hosted separately at the following addresses: hictkpy.readthedocs.io, paulsengroup.github.io/hictkR.

## 3 Results

### 3.1 Availability and Implementation

hictk is a header-only C++17 library accompanied by command-line tools to transparently operate on .hic and .cool files. hictk source code is available on GitHub at github.com/paulsengroup/hictk under the MIT license. Pre-built binaries for hictk can be obtained from bioconda (Linux and macOS). Docker images are available on GHCR.io and DockerHub. hictk can be built from source on Linux, macOS and Windows using a compiler toolchain supporting C++17, CMake v3.25+ and optionally Conan v2+. The C++ API can be used by other projects by including hictk source code using CMake or by installing hictk with Conan. hictkpy can be obtained through PyPI and bioconda or installed from source, while hictkR can be installed from source.

The CLI interface of hictk is built around facilities provided by the library component and is based on the very expressive CLI interface of cooler.

The interface consists of a single executable, named hictk with several subcommands (see Supplementary Table 2 from Supplementary Text 1).

Most hictk subcommands accept both Cooler and .hic files as input, and can output .hic, .cool, and .mcool files as needed. This is thanks to the hictk C++ library, which can natively read and write both Cooler and .hic files.

### 3.2 Benchmarks

We characterize hictk performance through a number of comparative benchmarks with .hic v8, v9 and .mcool files with close to 7 billion pairwise interactions (see Figs S2-S8; Supplementary Text 1 for more details). In summary, hictk is a near-drop-in, high-performance alternative to existing tools that can perform a variety of common operations on .hic and Cooler files. A thorough overview of the benchmark results, including benchmark graphs, is available in Supplementary Text 1 (Sections 4-11).

### 3.3 Current limitations

hictk supports the majority of operations enabled by cooler, straw and hic2cool, and features of the .hic and Cooler format specification. However, some aspects are not yet fully supported, including native operation on remote files which requires implementing a Virtual File Layer driver in the case of Cooler, and low level remote file operations in case of .hic files. Reading remote files can however already be achieved by mounting and accessing these on the local filesystem using e.g. FUSE. Support for reading or writing asymmetric cooler files is also lacking, albeit their usage is infrequent. Finally, hictk does not support reading or writing FRAG interactions from .hic files.

## Acknowledgements

We thank the ENCODE Consortium and the lab of Erez Lieberman Aiden for contributing with ENCODE data used as part of this study.

Unpublished genome assemblies and sequencing data for Schistocerca americana, Schistocerca cancellata and Schistocerca serialis cubense are used with permission from the DNA Zoo Consortium (www.dnazoo.org).

We thank the DNA Zoo and the Behavioral Plasticity Research Institute (BPRI) for providing the genome assemblies for Schistocerca americana, Schistocerca cancellata and Schistocerca serialis cubense.

## Funding

This work has been supported by the Norwegian Research Council (project 324137).

### Conflict of Interest

none declared.

## Notes

https://github.com/paulsengroup/hictk

https://github.com/paulsengroup/hictkpy

https://github.com/paulsengroup/hictkR

### Competing Interest Statement

The authors have declared no competing interest.

### Summary of Updates

The manuscript was updated to fit the "Bioinformatics Application Notes" format. As a result, all benchmarks have been moved to Supplementary Text 1. The benchmarks have been expanded and updated to use a more recent version of hictk. The manuscript now contains a reference to hictkR, a package which provides R bindings for hictk.

