## Supplementary Text 1 for "hictk: blazing fast toolkit to work with .hic and .cool files"

##### 1. Supplementary Table 1

**Supplementary Table 1**: Code snippets showing how to use hictk, hictkpy, and hictkR to read a subset of interactions from .hic, .mcool, and .cool files.

```cpp
#include <algorithm>
#include <cstdint>

#include <fmt/format.h>
#include <hictk/file.hpp>


hictk::File file{"interactions.hic", 10'000};
const auto selector = file.fetch("chr1:10,000,000-20,000,000");

// Write pixels to stdout
std::for_each(selector.template begin<std::int32_t>(),
              selector.template end<std::int32_t>(),
              [](const auto &pixel) {
                  fmt::print("{}\n", pixel);
              });
```

```python
import hictkpy as htk

import sys


file = htk.File("interactions.mcool", resolution=10_000)
selector = file.fetch("chr1:10,000,000-20,000,000")
df = selector.to_df()

# Write pixels to stdout
df.to_csv(sys.stdout, sep="\t", index=False, header=False)
```

```r
library(hictkR)


file <- "interactions.cool"
f <- File(file)

df <- fetch(f, "chr1:10,000,000-20,000,000", join=TRUE)

# Write pixels to stdout
write.table(df, sep="\t", row.names=FALSE, col.names=FALSE)
```

#### 2. Supplementary Table 2

**Supplementary Table 2**: List of subcommands available in hictk's CLI.

| **Subcommand** | **Description** |
|---|---|
| balance | Balance Hi-C matrices using ICE, SCALE, or VC. Balancing can be performed on genome-wide, inter-chromosomal, or intra-chromosomal interactions. Balancing is carried out using temporary files and without loading all interactions at once into memory. |
| convert | Convert matrices between .hic and Cooler formats. This includes converting all available resolutions and normalization methods. Conversion is performed in a streaming fashion, without loading all interactions into memory. |
| dump | Write interactions from .hic or Cooler files to the terminal in COO or Bedgraph2 format. hictk dump supports querying files for interactions overlapping specific regions using the --range and --range2 CLI options. Furthermore, hictk dump supports dumping balanced interactions, as well as expected and observed/expected interactions for .hic files. |
| fix-mcool | Fix corrupted .mcool files (e.g. files with corrupted file index). Corrupted files can be identified using hictk validate. hictk fix-mcool will attempt to re-balance the corrupted .mcool files using the same settings used to originally balance interactions. When the .mcool files lack the metadata with information on how the files were balanced, hictk will balance matrices using the default settings of hictk balance. |
| load | Build .cool and .hic files from interactions in 4DN DCIC pairs, validPairs, Bedgraph2, or COO format. Loading is performed using relatively little memory by taking advantage of temporary files. hictk load can generate single-resolution Cooler and .hic files. Multi-resolution Cooler and .hic files can be generated using hictk zoomify. |
| merge | Merge multiple Cooler or .hic files using the same reference assembly and resolution into a single file. Merging is performed by streaming interactions from Cooler or .hic files and merging pixels with the same coordinates. For Cooler files, this is performed without the use of temporary files, while for .hic files the use of temporary files is required to assign pixels to blocks of interactions without loading all interactions into memory at once. |
| rename-chromosomes | Rename chromosomes found in a Cooler file. Renaming can be performed in several ways:<br>- By adding the "chr" prefix to chromosomes (e.g. 1 -> chr1).<br>- By removing the "chr" prefix to chromosomes (e.g. chr1 -> 1).<br>- By providing a two-column TSV file mapping current chromosome names to new chromosome names. |
| validate | Validate Cooler and .hic files. Files are checked for various types of inconsistencies (e.g. for Cooler files, hictk checks whether the file is lacking mandatory attributes and fields, while for .hic files hictk ensures |

| | |
|---|---|
| | that the metadata for all chromosomes can be read without errors). When specifying the --validate-index CLI option, hictk will check the index of Cooler files to detect malformed indexes, such as those produced by cooler zoomify v0.8.3. The index corruption causes duplicate values to be returned for the affected row/column of interactions (meaning that a given bin pair is mapped to two, often different interaction counts). |
| zoomify | Convert single-resolution Cooler and .hic files to multi-resolution by coarsening. Interactions from the base resolution of the input .cool or .hic file is first copied to the output file. Then the base resolution is used to generate the coarser resolution, taking care to load as few pixels into memory as possible. Coarsening .cool files does not involve the use of temporary files, while coarsening .hic files does, as in the first pass, interactions are aggregated and assigned to the appropriate block of interactions, while in the second pass blocks of interactions are indexed and written to the final .hic file. |

### 3. Supplementary Table 3

**Supplementary Table 3**: File size, number of interactions and resolutions of .hic and .mcool files used for benchmarking.

| | hic v8 | hic v9 | Multi-resolution cooler |
|---|---|---|---|
| File size (GBs) | 176 | 77 | 53 |
| # of interactions | 6,974,747,832 | | |
| # of non-zero entries (10 bp resolution) | 5,873,705,034 | | 5,873,722,963 |
| Resolutions | 10 bp, 100 bp, 500 bp,<br>1 kbp, 5 kbp,<br>10 kbp, 25 kbp, 50 kbp,<br>100 kbp, 250 kbp, 500 kbp,<br>1 Mbp, 2.5 Mbp | | |

### 4. Benchmark system and data

Benchmarks were run on a system with the following specifications:

- AMD EPYC 7742 (2x 64 cores)
- 2048 GB (32x 64 GB, RDIMM DDR4 2933 MT/s eight-channel)
- RHEL 8.8 (Linux v4.18.0-425)

Input and output files were stored on a ramdisk to avoid IO bottlenecks.

The benchmark suite is written using Nextflow DSL 2 and was executed using Nextflow v23.04.1 (Di Tommaso *et al.*, 2017) limiting parallelism to at most 32 CPU cores to ensure results are not biased by high CPU load.

The following files were downloaded from ENCODE (ENCODE Project Consortium, 2012; Luo *et al.*, 2020; Hitz *et al.*, 2023) and UCSC FTP server and were used across benchmarks:

- ENCFF447ERX: pairs of interactions for GM12878 in text format (4DN-DCIC pairs) (identifiers.org/encode:ENCFF447ERX)
- ENCFF301CUL: list of Topologically Associating Domains (TADs) for GM12878 (identifiers.org/encode:ENCFF301CUL)
- hg38.chrom.sizes: list of chromosomes for hg38 (Lander *et al.*, 2001)

Interactions from ENCFF447ERX were used to generate interaction matrices in .hic v8, .hic v9 and .mcool formats using JuicerTools v1.22.01 (Durand *et al.*, 2016), HiCTools v3.30.00 (Durand *et al.*, 2016) and cooler v0.9.2 (Abdennur and Mirny, 2019) respectively. This dataset was selected as it is one of the largest dataset publicly available, with close to 7 billion interactions (see Supplementary Table 3). Each file contains 13 resolutions ranging from 10 bp to 2.5 Mbp. We decided to run our benchmarks with resolutions up to 10 bp, as this is the highest resolution available in .hic files released as part of ENCODE4 (www.encodeproject.org/pipelines/ENCPL839OAB) (ENCODE Project Consortium, 2012; Luo *et al.*, 2020; Hitz *et al.*, 2023). Each benchmark was repeated 3 times and median values and standard error were used for plotting.

The code used to generate the results presented in the next section is hosted on GitHub at github.com/paulsengroup/2023-hictk-paper and is archived on Zenodo at doi.org/10.5281/zenodo.10868277. Code used to generate matrices in .hic v8, .hic v9 and .mcool format can be found under preprocessing/ while code used for benchmarking is available inside the benchmarks/ folder. The repository README contains instructions on how to run the preprocessing and benchmark steps. Files used as input for benchmarks are automatically downloaded or generated by the preprocessing workflows and are also deposited at the Gene Expression Omnibus under identifier GSE242815 (identifiers.org/geo:GSE242815). Benchmark results and matrices used for visualization purposes throughout the paper are deposited on Zenodo at doi.org/10.5281/zenodo.10868775.

#### 5. Benchmarks: converting file formats

Converting .hic to .cool or .mcool files and vice-versa is achieved through the hictk convert subcommand with

```
1) hictk convert input.hic output.mcool
2) hictk convert input.mcool output.hic
```

The above command (1) will create a .mcool by copying all resolution and balancing vectors available in the .hic file given as input. Command (2) performs the same operations using .mcool files as input and outputting a .hic file. Supplementary Fig. 1. shows the result of converting .hic files to .mcool. Interaction frequencies are fully preserved by the conversion process. Correctness of the conversion is tested automatically by hictk's CI pipeline every time a pull request or commit is submitted to hictk's repository on GitHub.

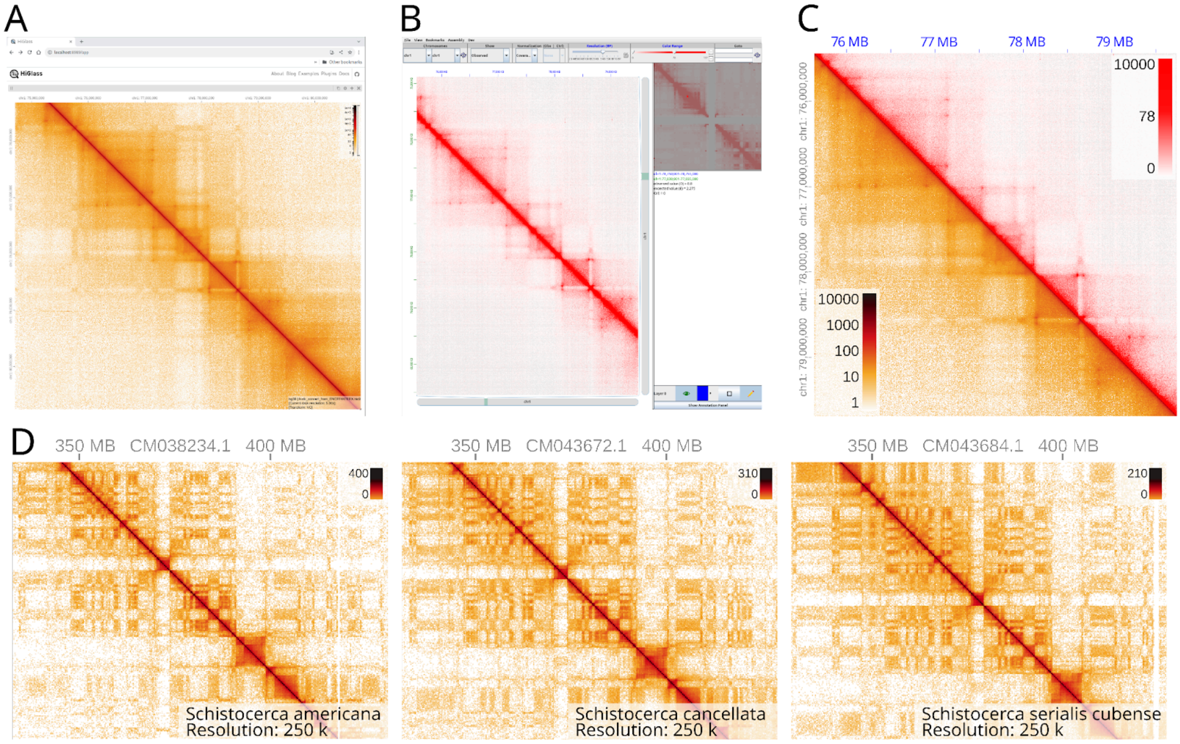


**Supplementary Fig. 1**: VC-normalized interaction matrices for hg38 chromosome 1 in .mcool and .hic visualized using HiGlass and JuiceboxGUI. Text elements too small to read are part of HiGlass and JuiceboxGUI and are not crucial to understand the figure. **A**: Interactions from a .mcool file visualized using HiGlass. **B**: Interactions from a .hic file visualized using JuiceboxGUI. **C**: Merged view of interactions visualized with HiGlass and JuiceboxGUI. **D**: Hi-C Interactions for three species of grasshopper from the DNA Zoo (Dudchenko *et al.*, 2017, 2018) visualized using HiGlass after converting matrices from .hic to .mcool using hictk convert.

We assess the performance of hictk convert by comparing the time required for converting files from .hic to .mcool format using hic2cool and hictk convert. For hictk we show the performance when converting from .hic v8 and v9 files. For hic2cool we only benchmark the performance when converting .hic v8 files, as this is the most recent standard supported by hic2cool.

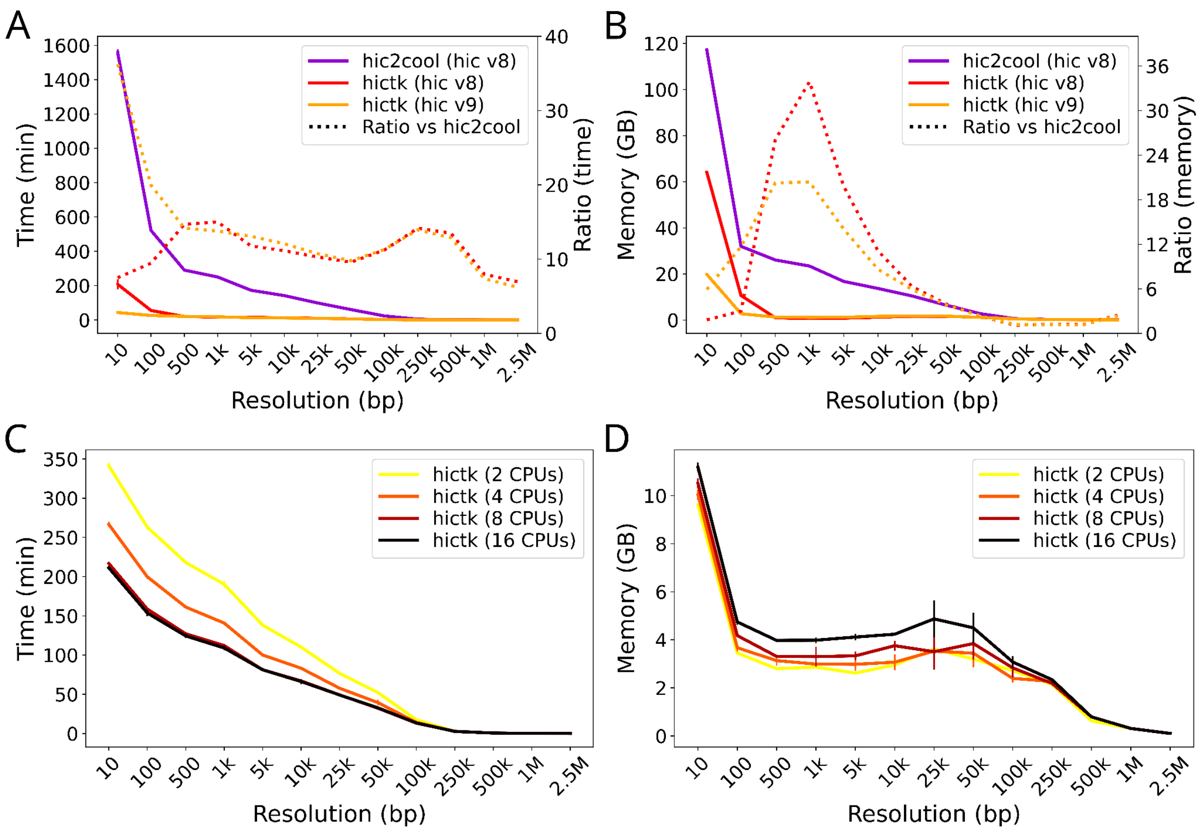


**Supplementary Fig. 2**: Benchmark results comparing the performance of hictk convert and hic2cool when converting .hic files to .cool format for different resolutions of the Hi-C dataset. Each combination of file format and tool is colored differently. Dotted lines show the relative improvement in performance and memory usage of hictk compared to hic2cool. Error bars show the standard error of time measurements taken across multiple runs of the same benchmarks **A**: Time required to convert a .hic file to .cool format. The Y axis on the right shows the performance improvement achieved when running hictk compared to hic2cool. **B**: Peak memory usage during conversion from .hic to .cool format. The right Y axis shows reduction of memory requirements when using hictk compared to hic2cool. **C**: Time required to convert a .cool file to a single-resolution .hic file. **D**: Peak memory usage during conversion from .cool to single-resolution .hic.

As seen in Supplementary Fig. 2A, when converting .hic v8 files, hictk is 7-15 times faster than hic2cool across all tested resolutions. It also maintains a comparable memory footprint at coarse resolution and notably consumes significantly less memory at finer resolutions (only 1/34 at 1000 bp resolution) (Supplementary Fig. 2B). When handling .hic v9 files, the disparity in performance and memory utilization becomes even more pronounced at higher resolutions. hictk outperforms hic2cool by a factor of 36 at a resolution of 10 bp, while consuming only one-sixth of the memory. At resolutions above 500 bp, hictk converts .hic v8 and v9 with a similar performance. At finer resolutions, however, performance is generally better for .hic v9 files. This performance difference is due to the new indexing strategy used in .hic v9 files, which leads to more compact indexes and interaction blocks that are less sparse and can be decompressed more efficiently.

hictk convert can also convert matrices from .mcool to .hic v9.

```
hictk convert input.mcool output.hic
```

This will generate a .hic file containing all resolutions found in the .mcool file. If balancing weights are available in the .mcool, those will be copied in the output .hic.

Conversion is achieved by assigning interactions to the interaction blocks that underlie .hic files. This is achieved through the use of temporary files. Conversion takes up to 5 hours 45 minutes when using two CPU cores (one core for input operations and the other for output operations) and up to 3h 32 minutes when using 16 CPU cores, where two cores are used for input/output operations, while the rest are used to compress blocks of interactions in parallel. Notably, the performance improvement obtained by using 16 instead of 8 CPU cores is negligible, suggesting that in the case of our test dataset, the compression pipeline can be saturated using 8 CPU cores when converting files containing high resolutions. To the best of our knowledge, hictk is the first tool to enable easy conversion from .mcool to .hic format.

We conclude that hictk convert is a drop-in replacement for hic2cool that is much more efficient at converting .hic files to .mcool format. Furthermore, hictk convert can be used to greatly simplify converting .mcool files to .hic format.

#### 6. Benchmarks: reading interactions

Reading interactions from .hic and .cool files is done with the hictk dump subcommand:

```
hictk dump input.hic --resolution 1000
hictk dump input.cool
hictk dump input.mcool::/resolutions/1000
```

This will write to stdout genome-wide interactions sorted by genomic coordinates in COO format (row, column, interaction count). Option --join can be used to output interactions in bedgraph2 format (chrom1, start1, end1, chrom2, start2, end2, interaction count). When available, balancing weights can be applied to interactions by passing the weight name to option --balance.

To fetch interactions overlapping a region of interest options --range and --range2 can be used:

```
hictk dump input.cool --range chr1
hictk dump input.cool --range chr1:0-10,000,000 --range2 chr2:0-5,000,000
```

hictk dump also supports querying the provided files for other data types and properties, such as chromosome table, bin table, as well as available normalizations, resolutions, and cells.

Next, we compare the performance of reading interactions from .hic8, .hic9 and .mcool files using cooler, straw and hictk. We measure the performance of dumping genome-wide as well as single chromosome interactions. Measurements for dumping genome-wide interactions using straw are missing as this operation is not supported by straw. Contrary to hictk and cooler, straw returns interactions sorted by block ID. Interactions are thus not guaranteed to be sorted by genomic coordinates. For this reason we also show the performance of a version of straw modified by us to sort interactions by genomic coordinates before returning them to the user (see Supplementary Text 1).

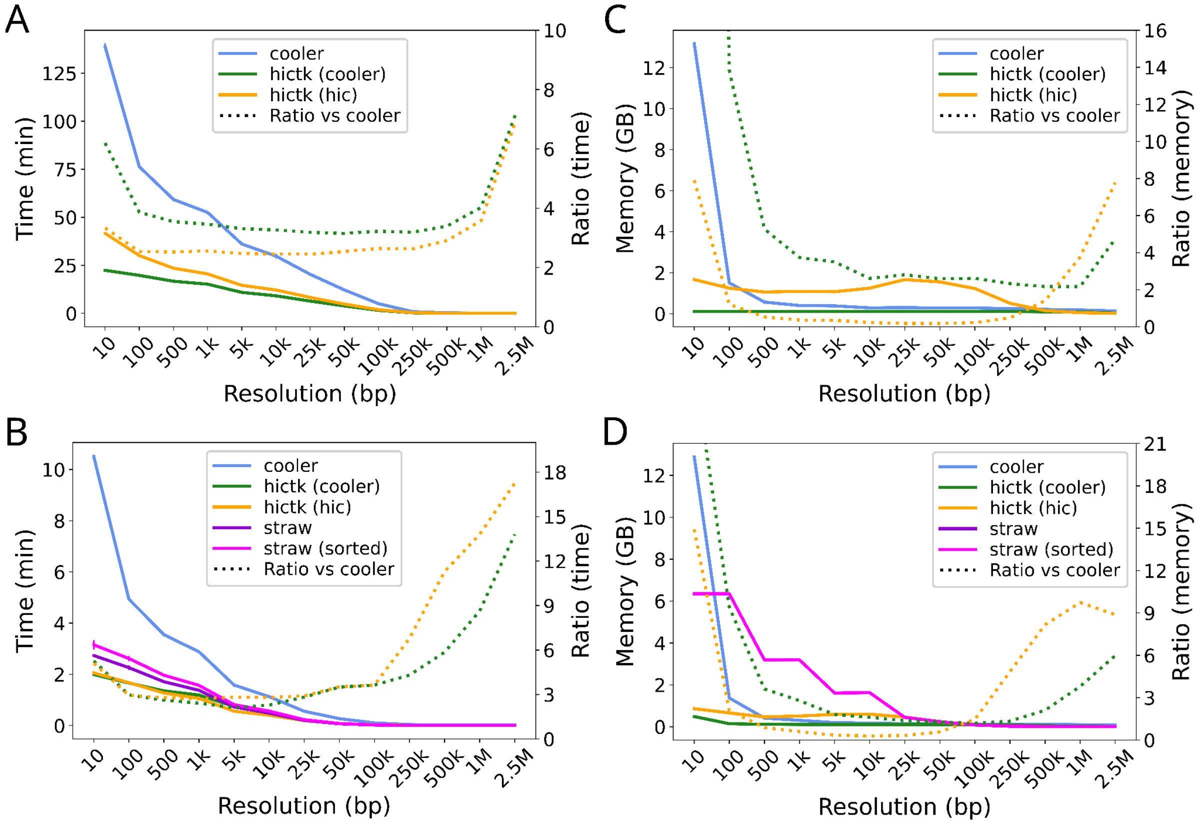


**Supplementary Fig. 3**: Benchmark results comparing the performance of hictk dump, cooler dump and straw for different resolutions of the Hi-C dataset. Dotted lines show the relative improvement in performance and memory usage of hictk compared to cooler. **A**: Time required to dump genome-wide interactions. The second vertical axis (right) shows the relative performance improvement achieved when running hictk compared to cooler. **B**: Time required to dump interactions for chromosome 1. The second vertical axis (right) shows the performance improvement achieved when running hictk compared to cooler. **C**: Peak memory usage while dumping genome-wide interactions. The second vertical axis (right) shows reduction of memory requirements when using hictk compared to cooler. **D**: Peak memory usage while dumping interactions for chromosome 1. The second vertical axis (right) shows reduction of memory requirements when using hictk compared to cooler. Note that the curve for “straw” appears to be missing because “straw” and “straw (sorted)” require exactly the same amount of memory.

Our benchmark shows that hictk is 3 to 18 times faster than cooler and 1.35 times faster than straw for dumping genome wide (Supplementary Fig. 3A) and chromosome 1 (Supplementary Fig. 3B) interactions. Compared to straw, hictk uses significantly less memory across all tested resolutions regardless of file format (Supplementary Fig. 3C and D). This is because hictk streams interactions using iterators, while straw buffers all interactions in memory before returning them to the user. The comparison with cooler memory requirements is more nuanced: when dumping interactions from .hic files, hictk uses more memory than cooler for resolutions greater than 1 kbp and approximately the same or less for higher resolutions; when dumping interactions from .cool files, hictk always requires less memory than cooler (Supplementary Fig. 3C and D).

Dumping interactions from .hic files with hictk is slightly slower than performing the same operation on .cool files (Supplementary Fig. 3A and 3B), as in this case interactions are stored separately for each pair of chromosomes, and hictk has to stitch together interaction

for pairs of chromosomes in the right order before printing them to stdout. This is also the reason why hictk requires more memory to dump genome-wide interactions from .hic files than from .cool files (Supplementary Fig. 3C and D). Dumping genome-wide interactions from .cool files is trivial, as interactions are already stored in a single vector sorted by genomic coordinates: this allows us to dump interactions without having to load the file index, which is why hictk uses very little memory to perform this operation (less than 15 MBs at 10 bp resolution) (Supplementary Fig. 3C).
In conclusion, hictk dump is significantly faster than available tools when dumping interactions for the entire genome or a single chromosome. Furthermore, hictk dump is the only tool to require less than 2 GBs of memory across all resolutions.

### 7. Benchmarks: creating .cool and .hic files from interactions in text format

Interactions in text format can be loaded into a .cool or single-resolution .hic file using hictk load. Supported formats are 4DN-DCIC pairs, validPairs, bedgraph2 and COO.

```
hictk load -f bg2 --bin-size 1000 chrom.sizes out.cool < interactions.txt
hictk load -f bg2 --bin-size 1000 chrom.sizes out.hic < interactions.txt
```

Interactions do not necessarily need to be sorted. However, if they are, the --assume-sorted flag can be used to efficiently load interactions in a single pass when creating .cool files. The flag is ignored when creating .hic files.

Next we benchmark the performance of creating .cool and single-resolution .hic files from interactions in text format using cooler cload, HiCTools pre, and hictk load. The benchmark consists of ingesting interactions in 4DN-DCIC pairs format into a .cool and .hic file. When creating .cool files, both cooler and hictk were run assuming interactions are in an unspecified order using a chunk size of 50,000,000. When creating .hic files, both HiCTools and hictk were allowed to use up to 16 CPU cores.

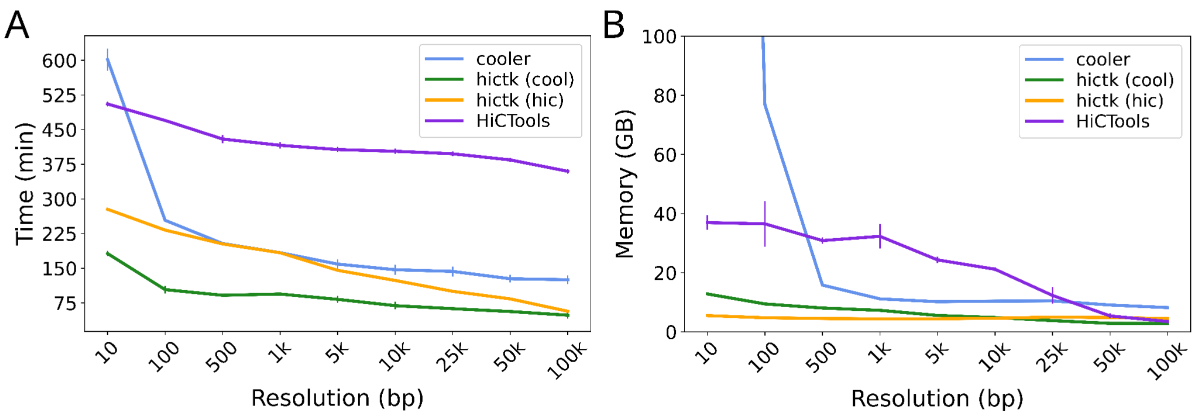


**Supplementary Fig. 4**: Benchmark result comparing the performance of hictk load, HiCTools pre, and cooler cload when creating .cool and .hic files at different resolutions. Error bars show the standard error of time measurements taken across multiple runs of the same benchmarks. **A**: Time required to ingest interactions at different resolutions. **B**: Peak memory usage while ingesting interactions at different resolutions.

To generate a .cool file at 10 bp resolution, hictk outperforms cooler by a factor of 2-3.5 ( Supplementary Fig. 4A). Memory requirements of hictk are also modest compared to those of cooler, necessitating 7 and 13 GBs of memory at 1 kbp and 10 bp, respectively, while cooler requires 11 and 760 GBs to process the same resolutions (Supplementary Fig. 4B).
Both hictk and cooler ingest interactions in two stages. In the first stage a fixed number of interactions are buffered and sorted in memory. Sorted interactions are then written to temporary files. Once all interactions have been processed, intermediate files are merged into the output cooler. This second stage is the most memory intensive in both applications. hictk uses a k-way merge algorithm using pairs of iterators and a priority queue which can merge k-files buffering exactly k-pixels in memory. In practice, for performance reasons, hictk buffers more than one pixel per file, but the number is low enough to require in the order of 1-100 MBs of memory per intermediate file.
When generating .hic files, hictk is 2-6 times faster than HiCTools, requiring approximately the same amount of memory when creating .hic and 50 and 100 kbp, and up to 8 times less when generating .hic files at finer resolutions.
We conclude that hictk load is a valid alternative to cooler cload and HiCTools across a wide range of resolutions, both in terms of processing speed and memory requirements. Furthermore, hictk requires the same parameters and flags to bin pair-wise interactions, regardless of the output file format, greatly simplifying generating .hic and .cool files, as it does not require interactions to be sorted in any particular order.

#### 8. Benchmarks: creating multi-resolution coolers

Single-resolution cooler and .hic files can be converted to multi-resolution files using hictk zoomify.

```
hictk zoomify input.cool output.mcool
hictk zoomify input.hic output.hic
```

The conversion is achieved by iterative coarsening, where a low resolution matrix is generated from a high resolution one.

We compare the performance of cooler coarsen and hictk zoomify when generating low resolution .cool files from higher resolution files by coarsening. Contrary to cooler, hictk can also zoomify single-resolution .hic files, such as the ones generated with hictk load. cooler coarsen supports processing files using multiple CPU cores, and was tested using 1, 4 and 8 CPU cores. hictk zoomify only supports multi-threading when generating .hic files, and was thus tested using 1, 4, and 8 CPU cores in this case, and using a single core when zoomifying .cool files.

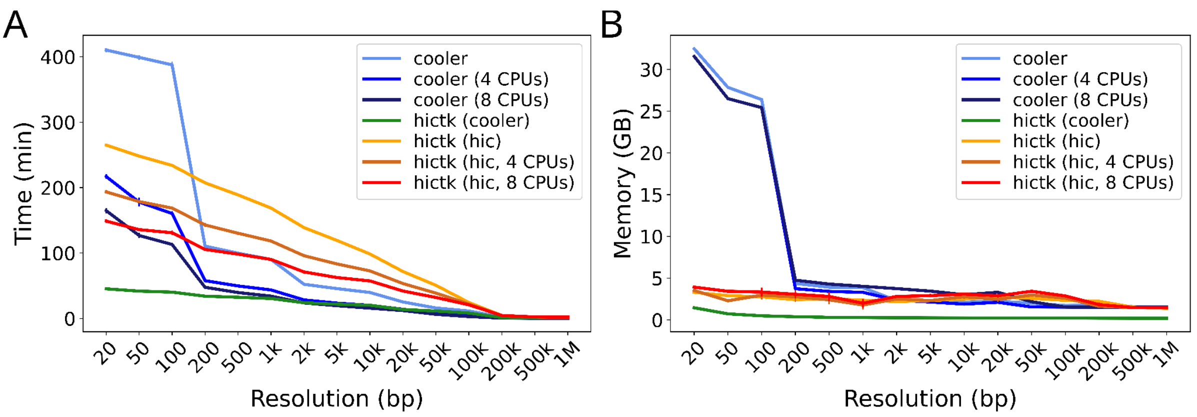


**Supplementary Fig. 5**: Benchmark results comparing the performance of hictk zoomify and cooler coarsen for different resolutions of the Hi-C dataset. The horizontal axis shows the resolution after coarsening. The greatest common divisor between the target resolution and resolutions listed in Table 1 was used as base resolution. Error bars show the standard error of time measurements taken across multiple runs of the same benchmarks. Dotted lines show the relative improvement in performance and memory usage of hictk compared to cooler using 1 CPU core. **A**: Time required to coarsen .cool files at different resolutions. **B**: Peak memory usage while coarsening .cool files.

When using a single CPU core to zoomify .cool files, hictk is faster than cooler across all tested resolutions. Yet, when cooler is allowed to use 4 or 8 CPU cores, hictk is relatively slower at low resolutions (greater than 20kbp). However, hictk is still 5 times faster when processing high resolutions (Supplementary Fig. 5A).

Generating a 20 bp .cool file from a 10 bp file takes less than 45 minutes with hictk and approximately 3.5 hours with cooler, even when utilizing 8 CPU cores (Supplementary Fig. 5A). Furthermore, hictk requires significantly less memory than cooler across all resolutions, using between 130 MBs and 1.5 GBs of RAM, while cooler requires at least 1.3 GBs and up to 35 GBs (Supplementary Fig. 5B).

Zoomifying .hic files requires more time than performing the same operation on .cool files because this requires two passes on the input data: one to assign interactions to interaction blocks, and another to merge and write blocks to .hic files. Zoomifying .hic files also requires more memory compared to .cool files, however memory requirements are relatively low when compared to those of cooler.

hictk takes advantage of a memory-efficient data structure to represent bin tables, which only requires storing chromosome names, lengths and the prefix sum of chromosome lengths, making memory requirements independent of resolution. This data structure allows for efficient mapping between bin identifiers (i.e. matrix rows and columns) and genomic coordinates. Given the above, coarsening in hictk amounts to the following operations: map interactions from the old coordinate system to the new one, then aggregate interactions mapping to the same pair of bins. Both operations are performed using iterators, buffering at most pixels overlapping one row at the lowest resolution. Interactions are aggregated using a hashmap and a priority queue. An implementation using binary-search trees was tested but found on average to be slower.

In conclusion, hictk zoomify can be used as a more efficient alternative to cooler coarsen and cooler zoomify to generate multi-resolution Cooler files by coarsening.

### 9. Benchmarks: balancing matrices using ICE

Hi-C matrices often are subjected to normalization by matrix balancing to remove experimental and technical biases. Both hictk and cooler implement matrix balancing through Iterative Correction and Eigenvector decomposition (ICE) (Imakaev *et al.*, 2012) hictk is capable of balancing both .hic and cooler files using the hictk balance subcommand:

```
hictk balance matrix.cool
hictk balance matrix.hic
hictk balance matrix.mcool
```

When provided with a multi-resolution .hic or cooler file, hictk will proceed to individually balance all available resolutions. hictk supports many parameters to tweak the balancing procedure, such as a --mode parameter to determine whether genome-wide, cis-only or trans-only interactions should be used for balancing. By default, hictk process interactions in chunks, avoiding loading the potentially large file into memory. However, when memory is not a constraint, users can specify the --in-memory flag to load all interactions into memory, significantly improving performance. Given that balancing a Hi-C matrix with ICE often involves traversing the same data hundreds of times, hictk first converts interactions from .hic or cooler format to an intermediate representation that is stored using temporary files. The file format used for the intermediate representation is only used internally by hictk and has been developed to enable efficient parallel iteration through Hi-C interactions, yielding a noticeable performance improvement compared to iterating through .hic or cooler files.

We compare the performance of hictk balance and cooler balance using two different benchmarks, one testing balancing performance using a fixed number of CPU cores (16) across many different resolutions (10-2.5M bp), and the other using a variable number of CPU cores (4-32) to balance a single resolution (500 bp). Both cooler and hictk were used to balance genome-wide interactions using up to 50 iterations. While hictk can balance both .hic and cooler files, we only benchmarked balancing of cooler files because hictk only traverses files in their original format once to convert them to the intermediate representation mentioned earlier. Thus, the expected performance difference when balancing .hic and cooler file is minimal.

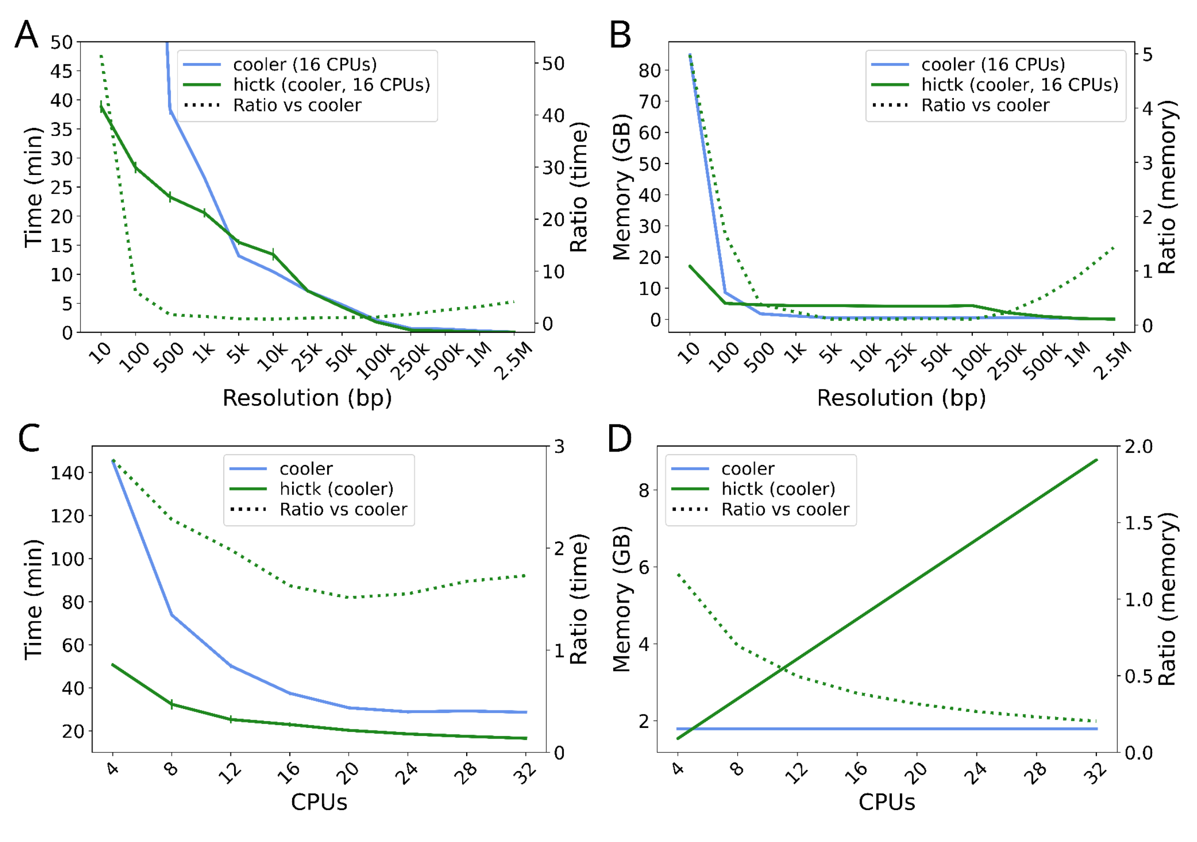


**Supplementary Fig. 6**: Benchmark results comparing the performance of hictk balance and cooler balance at different resolutions using 16 CPU cores and at 500 bp using a variable number of CPU cores. Dotted lines show the relative improvement in performance and memory usage of hictk compared to cooler. **A**: Time required to balance a .cool file at different resolutions. The second vertical axis (right) shows the performance improvement achieved when running hictk compared to cooler. **B**: Peak memory usage while balancing .cool files. The second vertical axis (right) shows reduction of memory requirements when using hictk compared to cooler. **C**: Time required to balance a .cool file at 500 bp resolution using a variable number of CPU cores. The second vertical axis (right) shows the performance improvement achieved when running hictk compared to cooler. **D**: Peak memory usage while balancing a .cool file at 500 bp resolution using a variable number of CPU cores. The second vertical axis (right) shows reduction of memory requirements when using hictk compared to cooler.

hictk and cooler exhibit similar performance when balancing a .cool file at 500 bp resolution using 16 CPU cores at resolutions greater than 5 kbp. However, as resolutions become finer and finer, a significant performance gap begins to appear, with hictk being 50 times faster than cooler at balancing matrices at 10 bp resolutions (Supplementary Fig. 6A). When it comes to memory usage, hictk tends to use more memory than cooler for resolutions greater than 100 bp. However the memory requirements remain quite modest, with hictk requiring less than 10 GBs of memory for all resolutions but the 10 bp resolution (Supplementary Fig. 6B). When balancing 500 bp .cool files using a variable number of CPU cores, hictk is approximately 2-3 times faster than cooler (Supplementary Fig. 6C). In this benchmark, hictk's memory usage is proportional to the number of CPU cores used for balancing. This is due to how parallel processing is implemented in hictk, where each worker thread is responsible for reading a chunk of interactions, decompressing it, and storing it into memory before processing. Because of this, hictk requires more memory than cooler when using more than 4 CPU cores for balancing. However, like for the previous benchmark, memory requirements remain relatively modest, with hictk requiring less than 10 GBs of memory to balance the test matrix using 32 CPU cores (Supplementary Fig. 6D).

#### 10. Benchmarks: fetching interactions overlapping pairs of TADs

Next, we sought to assess the performance of the library behind hictk by measuring the time required to fetch and sum interactions overlapping a pair of TADs. The benchmark consists of two parts, one testing the performance within chromosomes (cis queries) and the other between chromosomes (trans queries). Each part runs 5000 queries generated by random sampling with replacement on TADs from ENCFF301CUL (identifiers.org/encode:ENCFF301CUL). Cis queries are generated by using the same coordinates for both dimensions, while trans queries are generated by using coordinates from TADs mapping to different chromosomes. For each query we measure the time and memory required to fetch and sum interactions returned by the query.

This benchmark was designed to stress hictk's query engine similarly to what is done by applications fetching TAD-TAD interactions, such as applications and pipelines to call TAD cliques (Paulsen *et al.*, 2019; Rossini, 2023).

We compare the performance of hictk C++ API with cooler and hic-straw Python API using benchmark scripts fetch_and_sum_cooler.py, fetch_and_sum_straw.py and executable fetch_and_sum_hictk hosted on GitHub at github.com/paulsengroup/2023-hictk-paper

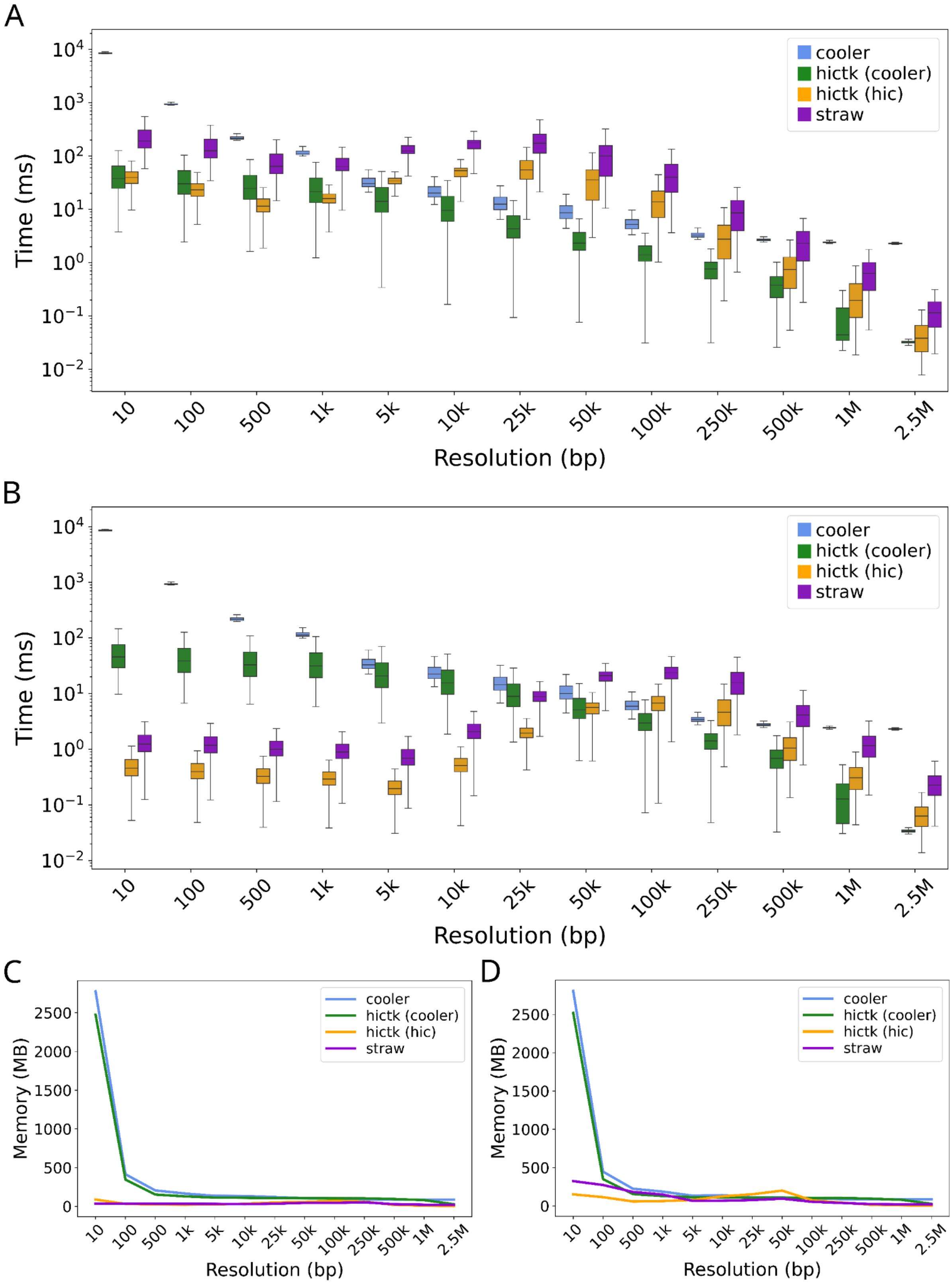


**Supplementary Fig. 7**: Benchmark results comparing performance when fetching interactions overlapping pairs of TADs. Queries were run for different resolutions of the Hi-C dataset. **A**: Time required to process queries for pairs of TADs located on the same chromosome (cis queries). **B**: Time required to process queries for pairs of TADs located on different chromosomes (trans queries). **C**: Peak memory usage when processing cis queries. **D**: Peak memory usage when processing trans queries.

For cis queries, the performance of hictk in reading .cool files stands out as the fastest among the tested tools (Supplementary Fig. 7A). It is faster for resolutions ranging from 5 kbp to 2.5 Mbp (with the exception of 1.0 Mbp). Notably, when working with resolutions of 1 kbp or higher, hictk's efficiency in reading .hic files becomes even more pronounced. In contrast, cooler lags behind as the slowest performer in both high and low-resolution scenarios, while straw is the slowest tool at intermediate resolutions. Overall, hictk is 3-6 times faster than cooler and straw at reading .cool and .hic files respectively. In the case of cooler, hictk is up to two orders of magnitude faster than cooler when processing interactions at very high resolution.

When processing trans queries, hictk is once again the fastest among the tested tools (Supplementary Fig. 7B). In general, hictk reading .cool file is faster when processing 50 kbp to 2.5 Mbp resolutions, while hictk reading .hic files becomes the fastest tool at resolutions higher than 50 kbp. Finally, at high resolutions, straw is faster than cooler, while at low resolution cooler is faster than straw. Overall, hictk is 2-6 times faster than straw at reading .hic files and 1.5-185 times faster than cooler at reading .cool files.

The memory requirements of all tested libraries when processing cis queries are similar for medium and low resolution, with hictk and straw consistently using slightly less memory than cooler (Supplementary Fig. 7C). At resolutions higher than 500 bp the memory required to process .cool files grows significantly for both cooler and hictk, as both libraries have to load a relatively large file index into memory in order to process queries. When processing .hic files, both straw and hictk require approximately the same amount of memory across all tested resolutions (Supplementary Fig. 7C). When processing trans queries, all libraries require approximately the same amount of memory to serve queries at 500 bp resolutions and lower (Supplementary Fig. 7D). At higher resolutions, the memory requirements for processing .cool files are once again significantly greater due to reading the file index into memory. However, hictk consistently requires slightly less memory than cooler to process .cool files (Supplementary Fig. 7D). The memory requirements of straw and hictk are similar for resolutions of 5 kbp and lower, while at higher resolutions the memory requirements of hictk are slightly lower (Supplementary Fig. 7D).

In conclusion, hictk is consistently faster than cooler and straw when reading .cool and .hic files respectively. In general, hictk achieves peak performance when reading from .hic and .cool files at high and low resolutions respectively. Finally, cooler performance is degraded at very high resolutions, with it being slower than other tools by up to 4 orders or magnitude, suggesting that under these circumstances the benchmark is testing code that has not been optimized for very high resolutions.

#### 11. Updating existing applications to use hictk

The original version of dejunlin/hicrep (Lin *et al.*, 2021) uses the cooler library to read files in .cool and .mcool format. We forked dejunlin/hicrep and updated the application to use hictkpy instead of cooler, enabling hicrep to read files in .hic, .cool and .mcool formats. The update involved adding 4 new lines of code, removing 59 and modifying 48 (see Supplementary Text 2).

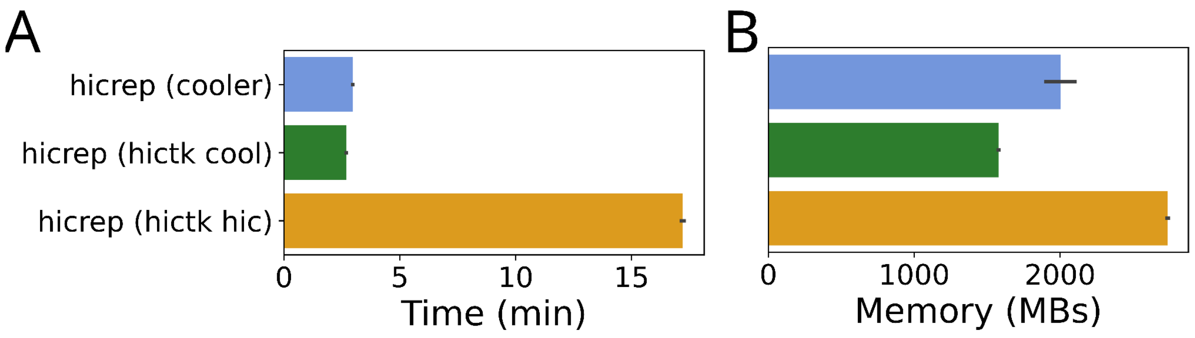


**Supplementary Fig. 8**: Benchmark results comparing the performance of hicrep using hictk and cooler as libraries to fetch interactions at 25 kbp resolution. Error bars show the standard error of time and memory measurements taken across multiple runs of the same benchmarks. **A**: Time required to compute the SCC for all human chromosomes. **B**: Peak memory usage while computing the SCC for all human chromosomes.

For reading .cool files, performance is similar using cooler or hictk (Supplementary Fig. 8A). Note, however, that computing the SCC on .hic files takes much longer than on .cool files due to a normalization step carried out by hicrep, which requires traversing .hic files in their entirety. This is not required when processing .cool files because the normalization factor is pre-computed and stored in the file. Memory requirements are slightly reduced by using hictk instead of cooler to process .cool files (Supplementary Fig. 8B). However, this is not the case when processing .hic files, where hictk requires more memory than cooler or hictk reading .cool files.
